# Anatomy and mechanics of tsetse fly blood feeding

**DOI:** 10.64898/2025.12.08.692967

**Authors:** Stephan Löwe, Laura Hauf, Elisabeth Meyer-Natus, Dianah Katiti, Dennis Petersen, Alexander Kovalev, Sebastian Büsse, Anna Steyer, Yoko Matsumura, Wolfgang Böhme, Daniel Masiga, Stanislav Gorb, Markus Engstler

**Affiliations:** Department of Cell and Developmental Biology, Biocentre, University of Würzburg, Germany; International Centre of Insect Physiology and Ecology, Nairobi, Kenya; Department of Functional Morphology and Biomechanics, Institute of Zoology, Kiel University, Germany; Electron Microscopy Core Unit, Department of Neurogenetics, Max Planck Institute for Multidisciplinary Sciences, Göttingen, Germany; Zoological Research Museum Alexander Koenig, Herpetology Section, Bonn, Germany

**Keywords:** tsetse fly, vertebrate host, skin, attachment, proboscis, probing forces, bloodmeal

## Abstract

African trypanosomiasis is a neglected tropical disease transmitted by tsetse flies (*Glossin*a spp.) in sub-Saharan Africa. The fly’s saliva carries parasitic unicellular trypanosomes, such as *Trypanosoma brucei*, leading to infections in humans, domestic animals, and wildlife. Tsetse flies feed on a broad spectrum of hosts, including humans, elephants, buffaloes, rhinos, hippos, turtles and monitor lizards. To understand how the insects can penetrate such diverse skin types, we detailed the anatomical structures involved in tsetse blood feeding, their mechanical properties and the forces exerted by the fly during probing. We found that the tsetse fly’s feeding apparatus does not rely on exceptionally unique structures or extraordinary forces. Instead, the tsetse fly has evolved subtle yet highly efficient adaptations, which include a labellum equipped with arrays of small teeth. When combined with a robust retraction movement of the proboscis, these structures create lesions on various types of skin, thus facilitating blood pool feeding on a broad host spectrum.

## Introduction

The World Health Organization (WHO) lists 20 neglected tropical diseases (NTDs) that affect about 1 billion people in the Global South (Aborode et al., 2022; Casulli, 2021; Engels & Zhou, 2020; World Health Organization, 2022). These infectious diseases are intrinsically linked to poverty (Hotez et al., 2006) and thrive where sanitation, clean water, and healthcare are lacking (Aborode et al., 2022; Bangert et al., 2017; Boisson et al., 2016).

Despite growing awareness and progress in controlling NTDs (Hotez et al., 2009; Molyneux et al., 2017; World Health Organization, 2024), post-COVID-19 effects and political instability in countries such as Sudan and the DRC have renewed threats to public health (Ehrenberg et al., 2021; Kavulikirwa, 2024; Manyazewal et al., 2024; World Health Organization, 2024), underscoring the fragility of recent progress and the need for sustained, coordinated action.

Several NTDs spread through insect vectors (Leitner et al., 2015), including the tsetse fly, responsible for transmitting African Trypanosomiases (Brun et al., 2010; P. Büscher et al., 2017). Tsetse flies are geographically restricted to a region in sub-Saharan Africa known as the ‘tsetse belt’ (Egeru et al., 2020; Meyer et al., 2016; Robinson et al., 1997), primarily because they are unable to traverse the vast expanse of the Sahara Desert (Ilemobade, 2009). Within this area, different *Glossina* species exhibit distinct ecological preferences, typically resulting in dominance within separate subregions (Attardo et al., 2019). All subspecies can be infected with trypanosomes and transmit them to vertebrate hosts during blood feeding.

Human African trypanosomiasis (HAT) is caused by *Trypanosoma brucei rhodesiense* and *T. b. gambiense*, while other trypanosome species, including *T. b. brucei*, *T. vivax*, *T. congolense*, *T. simiae, T. godfreyi*, and *T. suis* (Behrens, 1922; Rotureau & Van Den Abbeele, 2013; Steverding, 2017), give rise to animal diseases (Anene et al., 2001; Cattand et al., 2006; Gibson, 2007; Ugochukwu, 2009). Animal African trypanosomiasis (AAT), or Nagana, affects domestic animals such as cattle, sheep, goats, and horses across 40 countries (Griffin & Allonby, 1979; Ngeranwa et al., 1993; Singh & Idris, 2017; Spinage, 2012; Walden et al., 2014), severely hampering livestock productivity. This results in multi-billion-dollar economic losses (De Haan & Bekure, 1991; Feldmann et al., 2018; Ilemobade, 2009; McDermott & Coleman, 2001) that disproportionately impact rural and impoverished farming communities (Moonga & Chitambo, 2010; Odeniran et al., 2021; Wayo et al., 2017; Wilson et al., 1963).

Beyond humans and farm animals, tsetse flies feed on a remarkably broad range of wild vertebrate species (Bitome-Essono et al., 2017; Channumsin et al., 2021; Mwakasungula et al., 2022), many of which serve as important reservoir hosts for trypanosomes (Anderson et al., 2011; Dillmann & Townsend, 1979; Keymer, 1969; Kinghorn et al., 1913; Welburn et al., 2005). This opportunistic feeding behaviour extends to large, thick-skinned animals like Cape buffalos, hippopotamuses, rhinos, elephants, and lions, as well as reptiles such as crocodiles, snakes, and turtles (Bitome-Essono et al., 2017; Clausen et al., 1998; Weitz, 1963). In the Lake Victoria region of Kenya, riverine tsetse flies show a marked preference for feeding on *Varanus niloticus* monitor lizards, particularly in the early morning as they bask to elevate body temperature (Mohamed-Ahmed & Odulaja, 1997) (Kathiti et al., in preparation).

The extraordinary capacity to draw blood from such disparate hosts, successfully piercing through a lion’s dense fur as well as a lizard’s scales, points to a central knowledge gap in our understanding of tsetse feeding behaviour: How does this fly manage to efficiently exploit such a diverse host range? Does it possess specialized morphological or mechanical adaptations that confer this remarkable feeding flexibility?

To explore this, we examined the anatomy and mechanics of tsetse feeding in detail. We analysed how the tsetse fly attaches to its host by measuring the forces involved. In addition, we explored whether its mouthparts function as a ‘multifunctional tool’, equipped with adaptations that enable feeding across a broad host range. Finally, we assessed how the mechanical forces required for skin penetration vary, ranging from the tough hide of bovids to scaly lizard skin.

## Results

### The tsetse fly efficiently attaches to various surfaces

To initiate biting, the tsetse fly must first establish firm attachment to the host. The foot, or tarsus, plays a key role, as it makes direct contact with the host’s skin, fur, scales, or feathers. Using scanning electron microscopy, we examined the distinct structures of the tsetse tarsus. The sturdy claws (ungues) offer a secure grip on protruding features such as hair (Figure 1A- C, F). On the other hand, the flexible paired foot pads (pulvilli) are adapted for adhesion to relatively smooth surfaces like plant leaves, which the fly visits during resting hours (Figure 1A-C, F).

**Figure 1:**
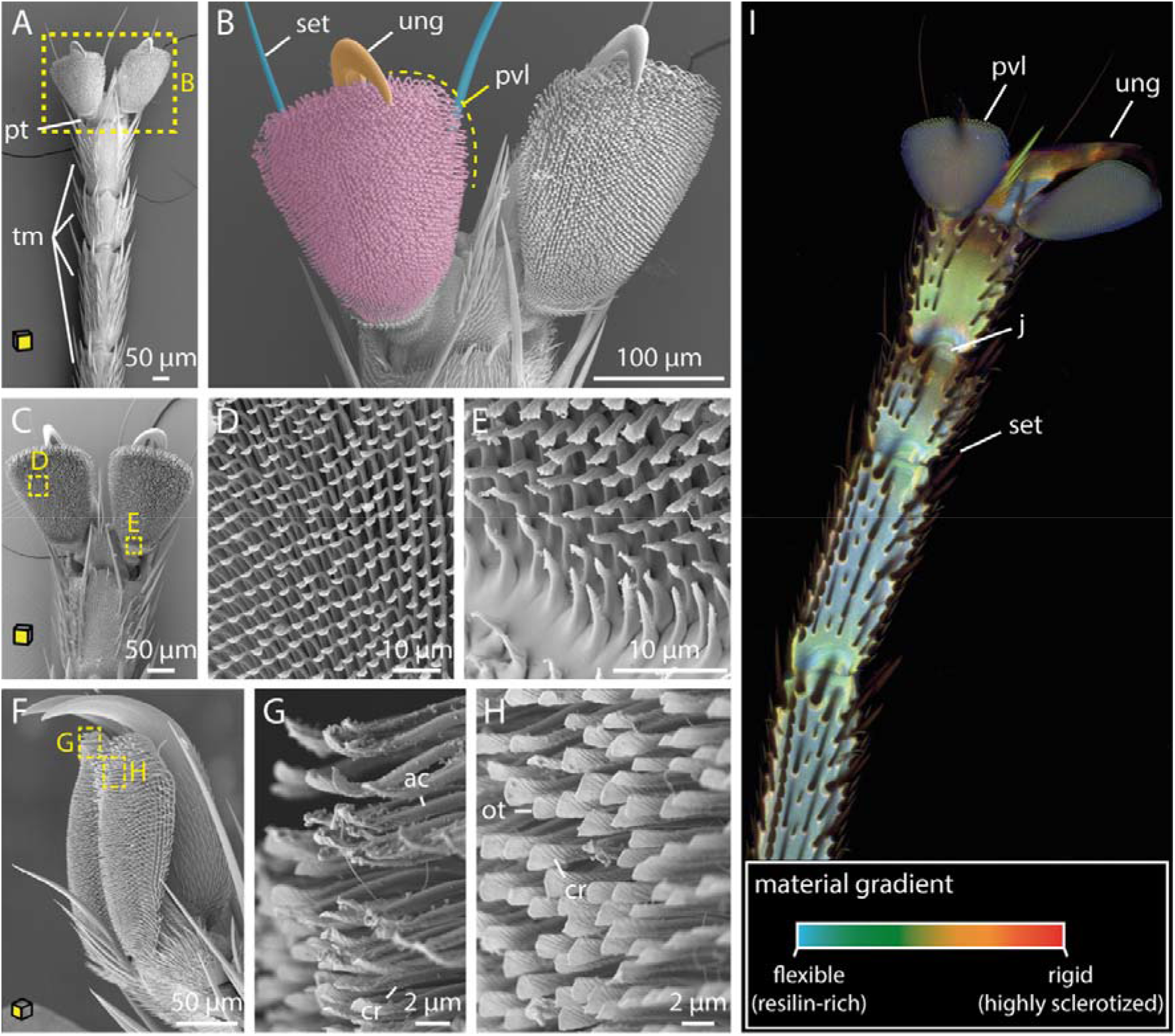
SEM and CLSM imaging of tsetse tarsi. **A-C:** Ventral views of the tarsi of the first (A, B), and second (C) leg pairs. **D-E:** Magnified views of the pulvillus from C**. F:** Lateral overview of the pulvillus and the ungues. **G-H:** Details of the pulvillus from F. Shown are the spatulas from a ventrolateral view at the rear of the pulvillus with signs of deformation (G), and further apart from the margin with no deformation (H). **I:** CLSM imaging representing compositional differences in the insect cuticle: flexible, resilin-rich regions appear blue, moderately stiff regions range from green to yellow, and rigid, highly sclerotized zones show up in red. Yellow planes on the cubes indicate the orientation of the tarsus attachment surfaces. **Abr**.: ac, acanthae; cr, crest; fr, furrow; j, joint; ot, oblate tip; pt, pretarsus; pvl, pulvillus; set, setae; sr, surface ridge; tm, tarsomeres; ung, ungues.

The pulvilli consist of densely packed fine cuticle outgrowths (acanthae), which increase in length and complexity from the proximal to the distal region (Figure 1C-H). The acanthae have a smooth and slender base that terminates in a flattened tip with submicron thickness, known as spatula (Figure 1D, E, G, H and Figure S1B). The dorsal side of the spatula features a proximal groove that follows its curvature, as well as several small knobs (Figure S1B). These knobs likely function to maintain physical separation between the acanthae (Haas & Gorb, 2004; Reinhardt et al., 2019). On the ventral side, a smooth surface transitions into a symmetrical crest-like structure (Figure 1G, H and Figure S1B). Notably, areas experiencing initial and most frequent surface contact exhibit signs of wear (Figure 1G).

The material properties of the tarsal structures were analysed by capturing exoskeleton autofluorescence using confocal laser scanning microscopy (CLSM) (Michels & Gorb, 2012). Resilin, a viscoelastic protein in the cuticle that is excited by UV light and emits blue fluorescence, plays a key role in mediating flexibility and elasticity (Michels et al., 2016; Niederegger & Gorb, 2003). In contrast, areas with extensive chitin-protein cross-linking (sclerotization) exhibit autofluorescence at longer, red-shifted wavelengths and impart stiffness (Michels & Gorb, 2012; Wootton, 2006).

In the tarsus, resilin distribution was found to be heterogeneous (Figure 1I). The central parts of the tarsomeres contain a higher concentration of resilin-dominated cuticle compared to the margins, except for areas around the joints. Conversely, the setae and ungues exhibit a stronger degree of sclerotization, suggesting higher rigidity. The pulvilli are resilin-dominated and thus likely exhibit high flexibility.

Overall, the tsetse fly tarsus exhibits neither anatomical nor mechanical specialization and is comparable to that of non-bloodsucking insects such as hoverflies or blowflies (Gorb, 1998; Gorb et al., 2001, 2012; Niederegger et al., 2002; Niederegger & Gorb, 2003).

To evaluate the functional performance of this generalist tarsal structure, we determined the attachment forces exerted by the tsetse fly on different surfaces (Figure 2A, Supplementary material 1, and Video 1). For this, male and female flies were placed on the spinning discs with varied surface roughness, ranging from smooth glass to textures with a granulation of 12 µm. For each fly, the rotational speed at the moment of contact loss was recorded, and the corresponding friction forces were calculated following the equations from Gorb et al., 2001. To normalize the attachment ability, the safety factor, which equals the total frictional force divided by the weight force of the fly, was determined (Figure 2E, F).

**Figure 2:**
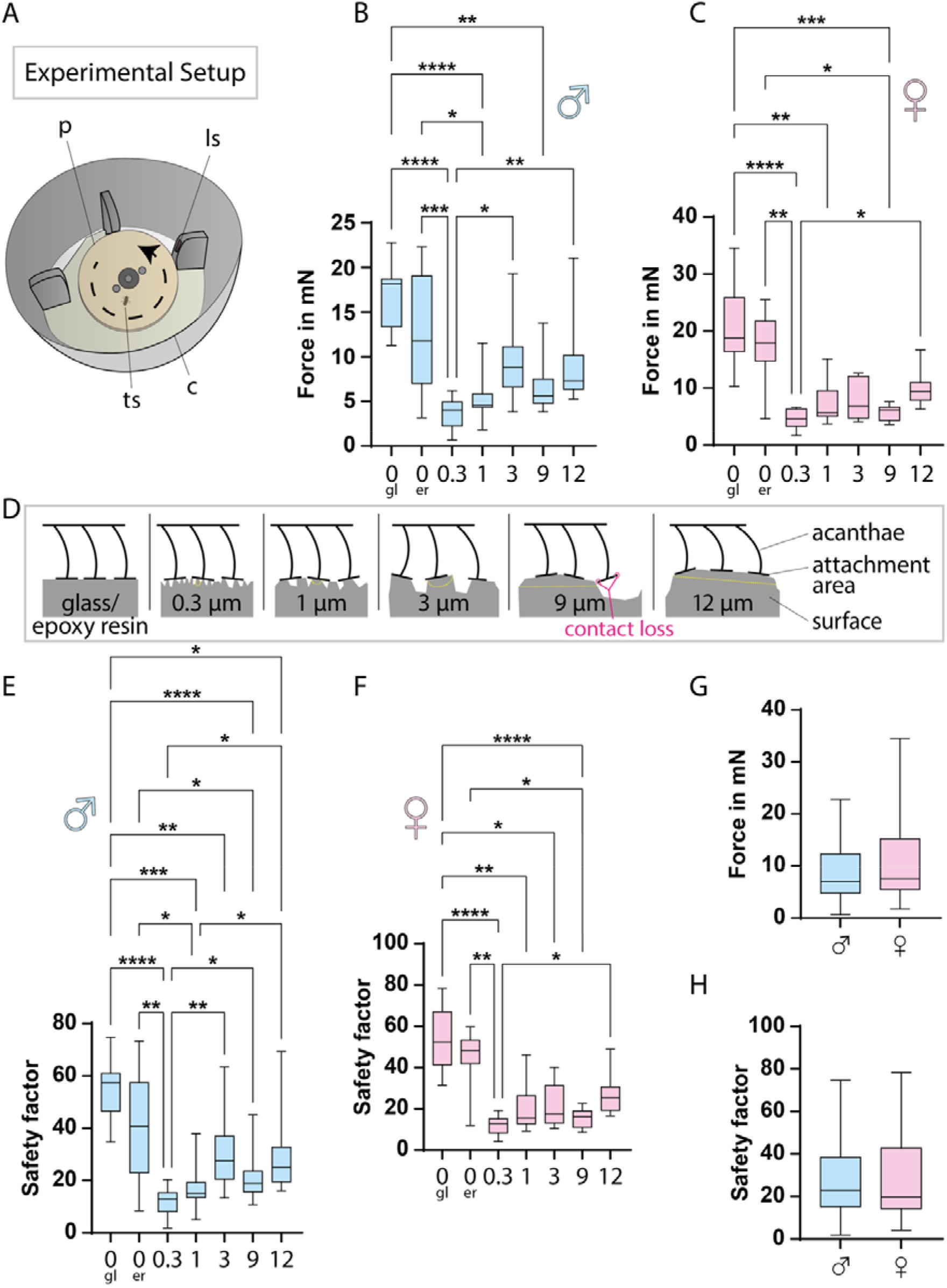
Tarsal attachment performance of tsetse flies expressed in friction force and safety factor. **A:** Schematic of the experimental setup for attachment measurements. The rotational plate, powered by a motor, accelerates until the fly loses grip, which is detected by a light sensor. **B-C:** Boxplots of friction forces at which male (B) and female (C) flies detached from different surfaces. Ten flies per sex were tested and for each surface, 3–14 replicates were averaged per fly. Graphs display these per-fly means, with boxes spanning the interquartile range, whiskers the full range, and a line marking the median. **D:** Illustration of how the acanthae interact with surfaces of different topographies. **E-F:** Safety factors for male (E) and female (F) flies on different surfaces. The safety factor, defined as the total friction force divided by weight force (mg, where m is body mass and g is gravitational acceleration), is a dimensionless quantity that enables cross-individual comparison. It was calculated for each replicate in B–C, and per-fly means were plotted. Boxplots are constructed as above. **G-H:** Boxplots of the friction forces (G) and safety factors (H) pooled across all substrates, compared between sexes, based on all per-fly means. Boxplots are constructed as above. Statistical significance in B, C, E, and D was determined using the Friedman test; *, p<0.05; **, p<0.001; ***, p<0.001; ****, p<0.0001. Comparisons between male and female flies in G and H were performed using the Mann-Whitney test, revealing no significant differences. All measured values, friction forces, and safety factors are provided in Supplementary material 1. **Abr**.: co, cover; gl, glass; er, epoxy resin (smooth); ls, light sensor; ps, surface plate; ts, tsetse fly.

The highest friction forces were measured on very smooth surfaces, such as glass or smooth epoxy resin (Figure 2B, C), which facilitate attachment of the fly with the majority of acanthae tips (Figure 2D). On these substrates, maximum forces and safety factors reached 34.5 mN and 78 for a female fly, and 22.8 mN and 75 for a male fly. Across all tested surfaces, median forces for female flies were 4.6 - 18.8 mN, corresponding to safety factors of ∼ 13 - 52. For male flies, median forces ranged from 4.0 - 18.2 mN, corresponding to safety factors of ∼ 13 - 57. Thus, there are no marked differences in the attachment capabilities of female and male tsetse flies (Figure 2G, H).

### The tsetse head and blood feeding apparatus

Having analysed the attachment structures of the tarsi, we next examined the tsetse head and feeding apparatus to explore adaptations for skin penetration and blood intake. While the front and lateral regions of the head are prominently occupied by large compound eyes (Figure 3A, B and Video 2), the ventral part is dominated by the mouthparts, consisting of the maxillary palps and the proboscis (Figure 3A-D, Figure S2A-C, and Video 2). The ventral section contains key digestive components, including the pharynx, which connects to the rest of the alimentary tract through the esophagus located in the thorax (Figure 3D, E and Figure S2C).

**Figure 3:**
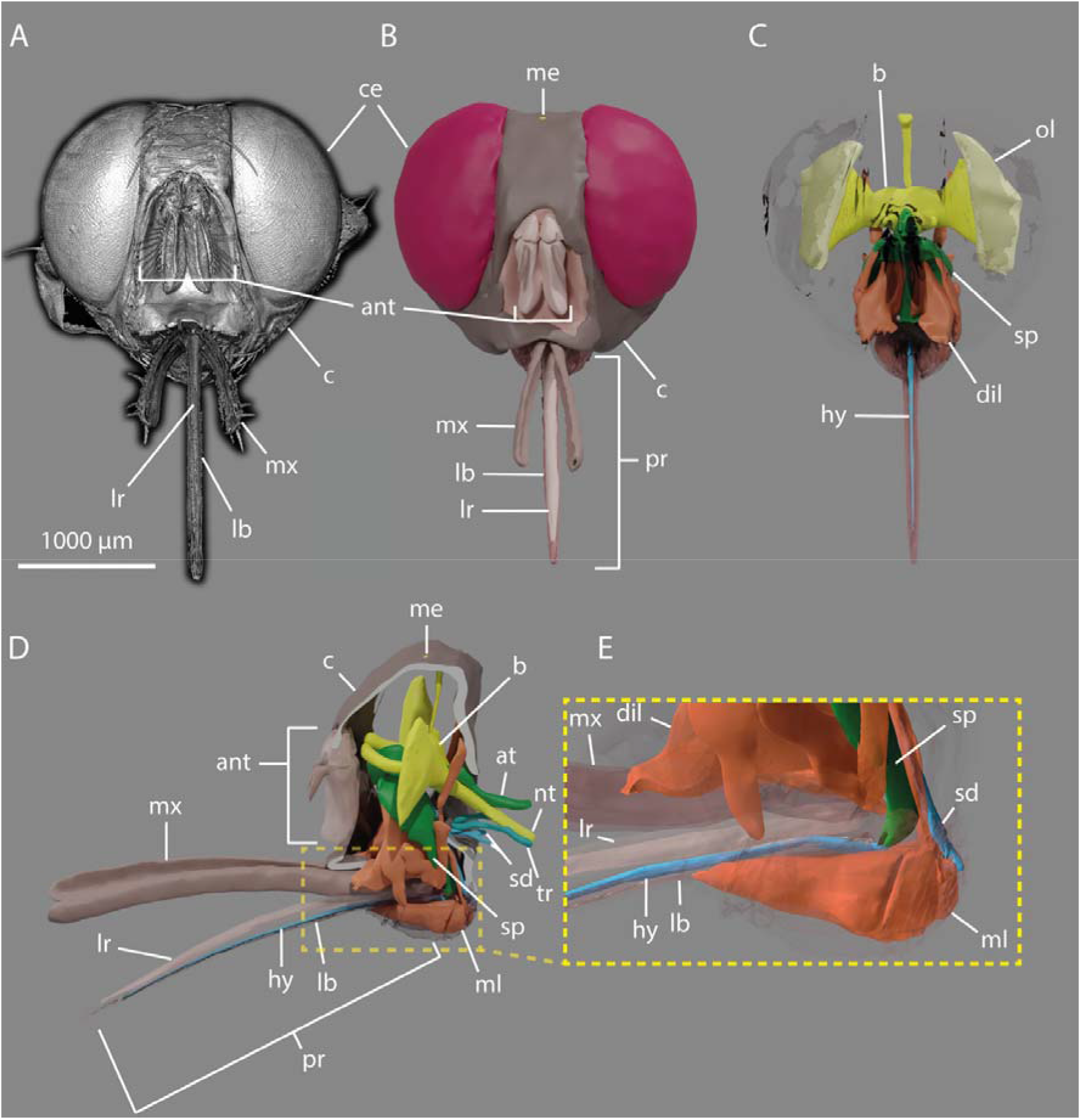
3D-reconstruction of the tsetse head. **A:** Frontal view rendering of a tsetse head derived from micro-computed tomography (µCT) data. Video 2 provided with this manuscript shows a 360° rotation of this 3D rendering. **B-C:** Frontal views of the 3D-reconstructed model from the same dataset, showing the external surface (B) and internal structures (C) **D:** Lateral view of C. **E:** Enlarged view of D emphasizing muscular structures associated with the proboscis and suction pump. **Abr.:** ant, antennas; at, alimentary tract; b, brain; c, caput; ce, compound eye; dil, pharyngeal dilator muscles; hy, hypopharynx; lb, labium; lr, labrum; me, median eye; ml, labial muscles; mx, maxillary palp; p, proboscis; ol, optical lobe; sd, salivary duct; sp, suction pump; tr, trachea; nt, thoracic nerve.

The maxillary palps serve as a protective structure to cover the proboscis, which is the central feature of the feeding apparatus (Figure 4A, D, E). The outer surface of the palps bears numerous robust setae of varying sizes (Figure 4A, D, E). The inner surface, in direct contact with the proboscis, is densely lined with fine, short setae (Figure 4E).

**Figure 4:**
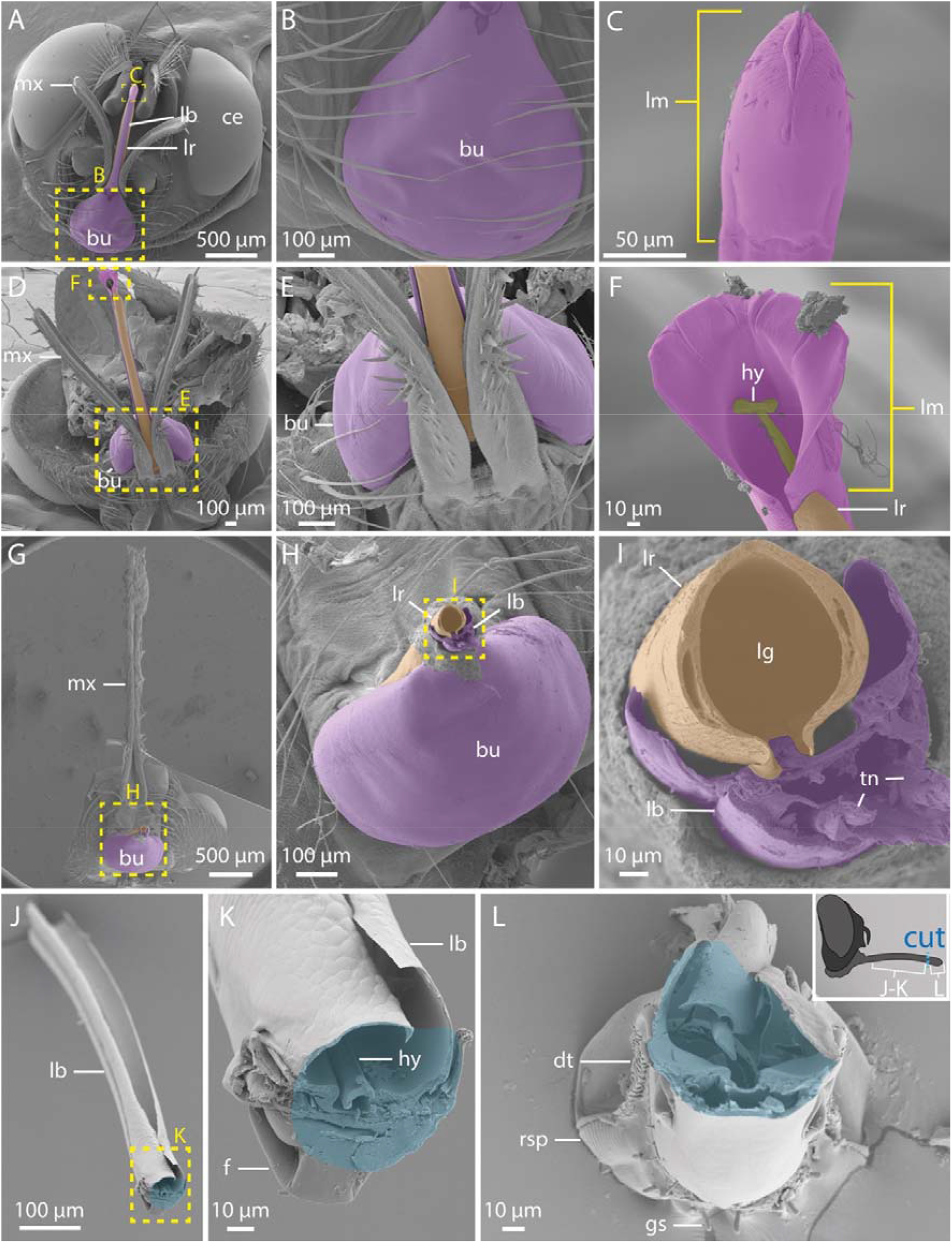
SEM micrographs of the tsetse proboscis and labellum joint details. **A:** Frontal view of the head showing the exposed ventral mouthparts**. B-C:** Enlarged views of A, focusing on the bulbus (B) and the closed labellum (C). **D:** Dorsal overview of the mouthparts. **E-F:** Magnifications of D, showing the dorsal base of the proboscis and maxillary palps (E), and the dorsal side of the opened labellum (F). **G-I:** Successive magnifications of a tsetse head after removal of the proboscis. The labrum displays hollow segments along its lateral walls (I). **J-L:** Views of a dissected proboscis. The schematic to the right indicates the separation site in blue, and the blue areas in the SEM images mark the corresponding cut surfaces where the parts were previously connected. Shown are the proximal portion (i.e. base to mid-length) of the proboscis as dorso-frontal overview (J) and a magnified detail of this highlighting the furca and labial gutter (K), as well as a view into the distal tip of the labellum (L). In an intact fly, the labrum would be positioned within the empty groove of the labium visible in J; however, it is absent in this dissected preparation. **Abr.:** bu, bulbus; ce, compound eyes; dt, dorsal teeth; f, furca; gs, gustatory sensilla; hy, hypopharynx; lb, labium; lg, labial gutter; lm, labellum; lr, labrum; mx, maxillary palps; rsp, rasping teeth; tn, tendons.

The proboscis itself is a functional unit. It consists of the labrum (upper lip) and the labium (lower lip) which jointly form a smooth, tube-like channel for blood uptake (Figures 3, S2, and 4A). Located in the ventral part of the head, the bulbus region of the proboscis anchors the labium (Figure 4A, B, D, E, G, H) and houses the labial muscles (Figures 3D, E, S2C, and S3D). Within the labium, tendons connect these muscles to the furca, a spring-like cuticular element (Figure 4K and Figure 5I). Coordinated muscle activity and tendon movement alter the configuration of the furca, enabling eversion of the labellum (Figure 4F and Figure 5C-G, I).

**Figure 5:**
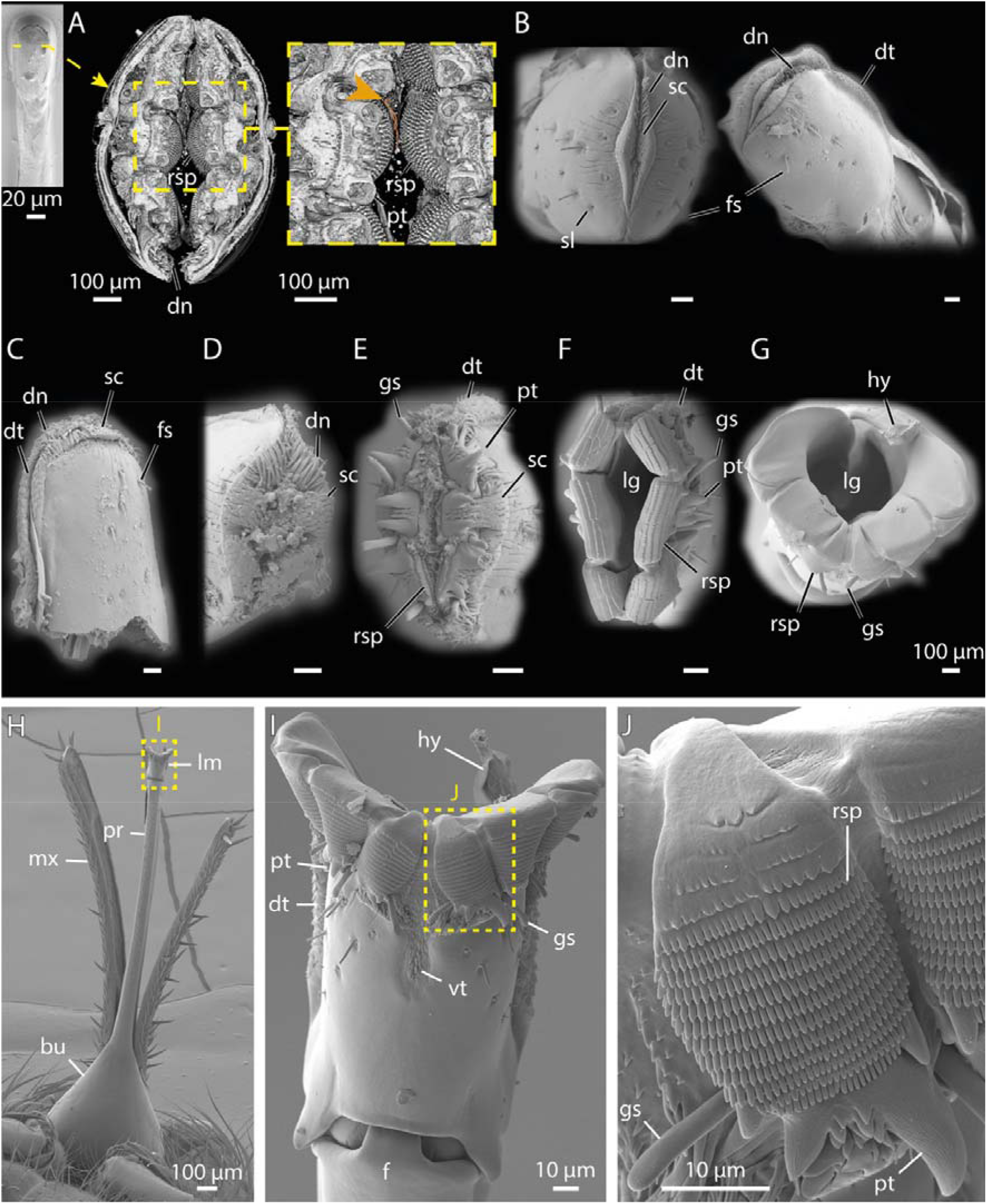
FIB-SEM and SEM imaging of the tsetse labellum and associated tooth structures. **A:** 3D reconstruction from FIB-SEM data (Video 3) showing the internal structure of the closed labellum in frontal view. The orange arrow indicates trypanosomes. A ventral view of the closed labellum is shown at the side for orientation. Video 3 provided with this manuscript shows a sequential progression through the labellum based on the FIB-SEM dataset. **B-G:** SEM micrographs of labella in progressively opened states; scale bars = 100 µm**. H:** Ventral overview of the proboscis with an open labellum**. I:** Close-up from H, highlighting exposed teeth-like structures. **J:** Enlarged view of I with focus on the rasping teeth. **Abr**.: bu, bulbus; dn, denticles; dt, dorsal teeth; fs, flagelliform sensilla; gs, gustatory sensilla; hy, hypopharynx; lg, labial gutter; lm, labellum; mx, maxillary palps; pr, proboscis; pt, prestomal tooth; rs, rasping teeth; sc, scales; sl, small sensilla; vt, ventral teeth.

At the proboscis tip, the labellum serves as the piercing component (Figure 4C, F and Figure 5). Its outer surface is equipped with fine sensilla. At the distal end of the labellum, a lip-like sclerite surrounded by numerous cuticular folds defines the opening region and mediates initial contact with the host tissue (Figure 5B).

Between the closed and fully open labellum, we observed multiple intermediate positions, each revealing different structures involved in chemical and mechanical perception, tissue laceration, tissue destruction, and tissue anchoring (Figure 5B-G). Scales and denticles covering the entire tip of the labellum cause minor cuts and abrasions in the host’s skin, resulting in shallow wounding (Figure 5A-E). Deeper lacerations are achieved by six large prestomal teeth arranged like flower petals around the labellum tip (Figure 5A, E, F, I, J and Figure S3B). These teeth alternately extend and retract to progressively deepen the wound. Additionally, rasping teeth located on the labial tip segments (Figure 5A, E, F, G, J, Figure S3B, C, and Video 3) bear numerous rows of microscopic cusps, further aiding in tissue disruption. When the labellum is closed, these structures fold inward, concealing them from external view (Figure 5A, B and Figure S3B, C). Thus, the anatomy and mechanics of the tsetse labellum resemble a saw that generates extended wounds for pool feeding of host blood.

The hypopharynx, a slender tube about 10 µm in diameter, is nestled within a slightly wider groove of the ventral part of the labium (Figure 4K and Figure S3E, F) and extends along the full length of the proboscis (Figure 3C, D). Its distal end reaches the anterior section of the labellum, where it forms finger-like protrusions (Figure S3B, C). Embedded within the hypopharynx is the salivary duct, which transports saliva from the proboscis to the anterior end of the labellum (Figure 3D, E).

The labium and labrum show high amounts of resilin-dominated cuticle (Figure 6A-D). In contrast, the labellum is more sclerotized, particularly at the prestomal teeth, rasping teeth, and across most of its surface (Figure 6A-D). Nonetheless, several flexible zones are present within the labellum, including the membranous areas surrounding the furca (Figure 6A-C), the lip-like sclerite at the labellum tip (Figure 6A), and the connective tissue around the rasping teeth (Figure 6D).

**Figure 6:**
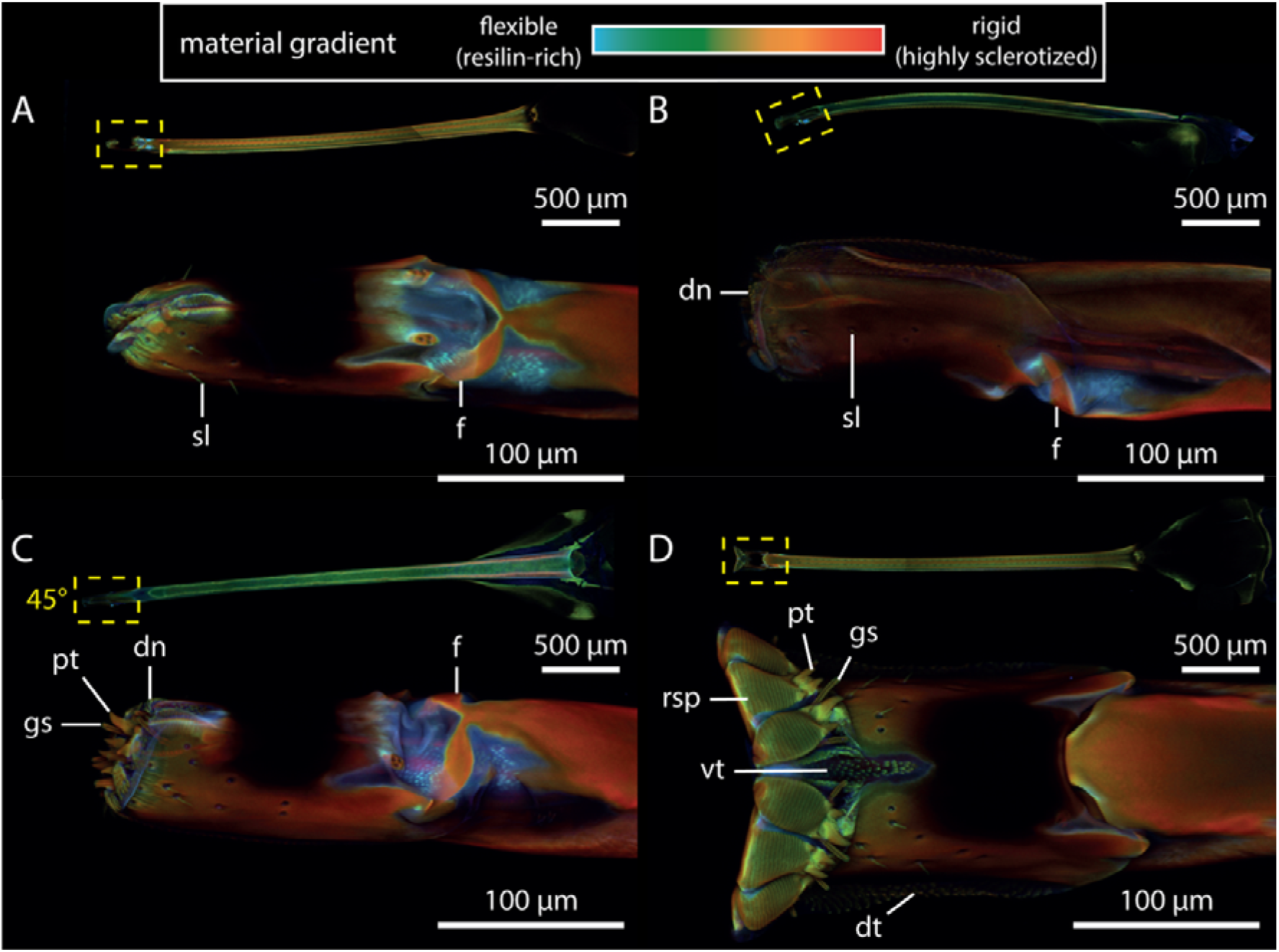
CLSM maximum intensity projections of tsetse probosces in different opening states, highlighting resilin distribution and sclerotization. The colour gradient represents compositional differences in the insect cuticle: flexible, resilin-rich regions appear blue, moderately stiff regions range from green to yellow, and rigid, highly sclerotized zones show up in red. **A:** Ventral view of the closed labellum, showing resilin-rich areas around the furca and the lip-like structure at the tip. **B:** Slightly opened labellum in lateral view, exposing denticles and the elevated position of the furca. **C:** Partially opened labellum in dorso-lateral view, exhibiting prestomal teeth and gustatory sensilla. **D:** Open labellum in ventral view, highlighting teeth-like structures. The full proboscis is visible at the top of each panel for orientation. In panel C, the perspective of the proboscis is rotated by 45° along its axis to show the magnified view of the labellum. **Abr**.: dn, denticles; dt, dorsal teeth; f, furca; gs, gustatory sensillum; pt, prestomal teeth; rsp, rasping tooth; sl, sensillum; vt, ventral teeth.

### Substrate interactions and force dynamics of the tsetse proboscis

Next, we analysed tsetse biting on a series of progressively more complex substrates. this included artificial substrates such as defined polydimethylsiloxane (PDMS) membranes and silicone feeding mats embedded with a grid of nylon threads (Figure S1A and Figure 7A-C), advanced human skin equivalents (Reuter et al., 2023), and *ex vivo* skin samples of human, cow, deer, and monitor lizard (Figure 7D-N).

**Figure 7:**
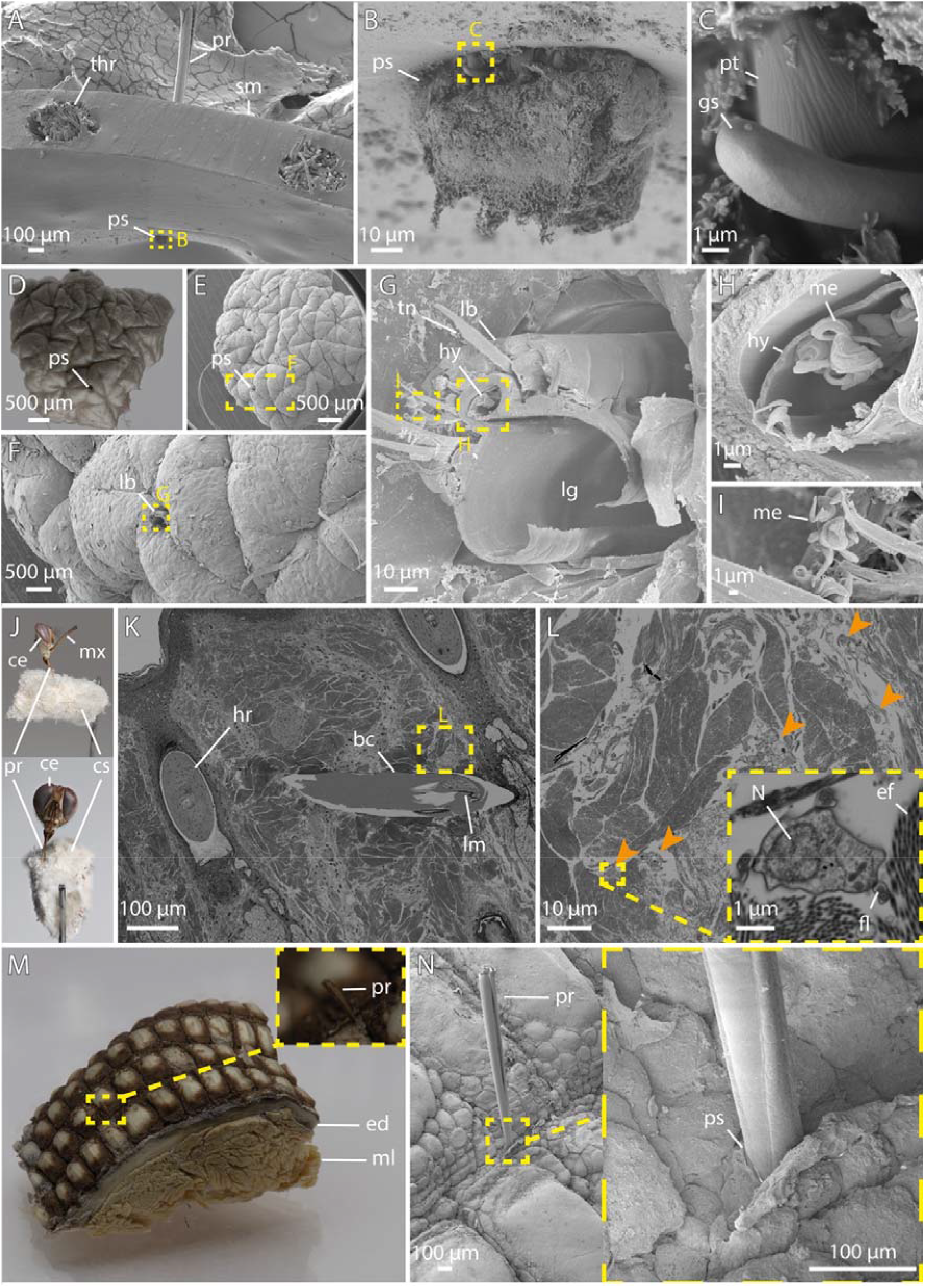
Macro and SEM imaging of tsetse proboscis interactions during feeding. **A-C:** Lateral views of a tsetse proboscis piercing through a silicone feeding mat. The labellum is partially protruding through the material (B), and a prestomal tooth is visible (C). **D-G:** A human skin explant showing a tsetse bite site, with remnants of the proboscis retained in the tissue (F, G). **H-I:** Higher magnifications of the proboscis from G, showing the hypopharynx filled with metacyclic trypanosomes (H), and expelled trypanosomes adhered to adjacent regions of the proboscis (I). **J:** A tsetse fly feeding on cow skin; the body was removed and the sample subsequently fixed. **K-L:** Ultra-thin section of resin-embedded cow skin. Shown is a proboscis within the biting channel (K). Successive magnifications reveal trypanosomes, indicated by orange arrows, in the surrounding tissue (L). **M-N:** Overview and SEM detail of *V. niloticus* lizard skin with a tsetse proboscis inserted between the scales. **Abr**.: bc, biting channel; ce, compound eye; cs, cow tissue; ed, epidermis; fl, flagella; gs, gustatory sensillum; hr, hair root; hy, hypopharynx; lb, labium, lg, labial gutter; lr, labrum; me, metacyclic trypanosome; ml, muscle layer; mx, maxillary palp; N, nucleus; pr, proboscis; pt, prestomal tooth; ps, penetration site; sm, silicone membrane; thr, thread; tn, tendon.

Visualizations of the tsetse bite, including both standard and high-resolution imaging, provided insights into how the proboscis interacts with tissue during feeding (Figure 7). Despite the prominent outward-pointing teeth (Figure 7B, C), the proboscis creates a remarkably smooth-edged channel (Figure 7D-G, K). Notably, trypanosomes were present in the fly’s hypopharynx (Figure 7G, H), as well as the tissue surrounding the bite site (Figure 7L). This rapid spread demonstrates the parasite’s high motility and efficient navigation even within dense and confined environments (Reuter et al., 2023; Schuster et al., 2017).

While probing sites on human and cow skin appeared unrestricted, tsetse bites on monitor lizard skin seemed confined to areas between the larger scales, where smaller, presumably softer scales allow penetration (Figure 7M, N).

A tsetse bite can be divided into four main stages: initial penetration, probing, blood uptake, and final proboscis retraction. To quantify and compare the forces exerted on different substrates, flies were induced to bite samples mounted on a 3D-printed platform connected to a force sensor.

All forces remained in the low mN range (Figure 8A, C-F, Figure S4, and Supplementary material 2). Penetration forces differed markedly across substrate types, reaching their peak on artificial materials with median values of 1.64 mN for PDMS and 1.63 mN for the silicone feeding mat (Figure 8C, D). Penetration forces on animal skins were intermediate, with median values of 1.34 mN for cow skin (which showed the highest single force of 3.1 mN), 1.13 mN for deer skin, and 1.37 - 1.38 mN for monitor lizard skin from both dorsal and tail regions (Figure 8C, D). Human skin substrates showed the lowest penetration forces, with dermal equivalents requiring less force (0.67 mN) than full-thickness skin equivalents (0.97 mN), reflecting the additional mechanical resistance of the epidermal layer absent in dermal-only constructs. Native skin explants fell intermediate at 0.85 mN (Figure 8C, D).

**Figure 8:**
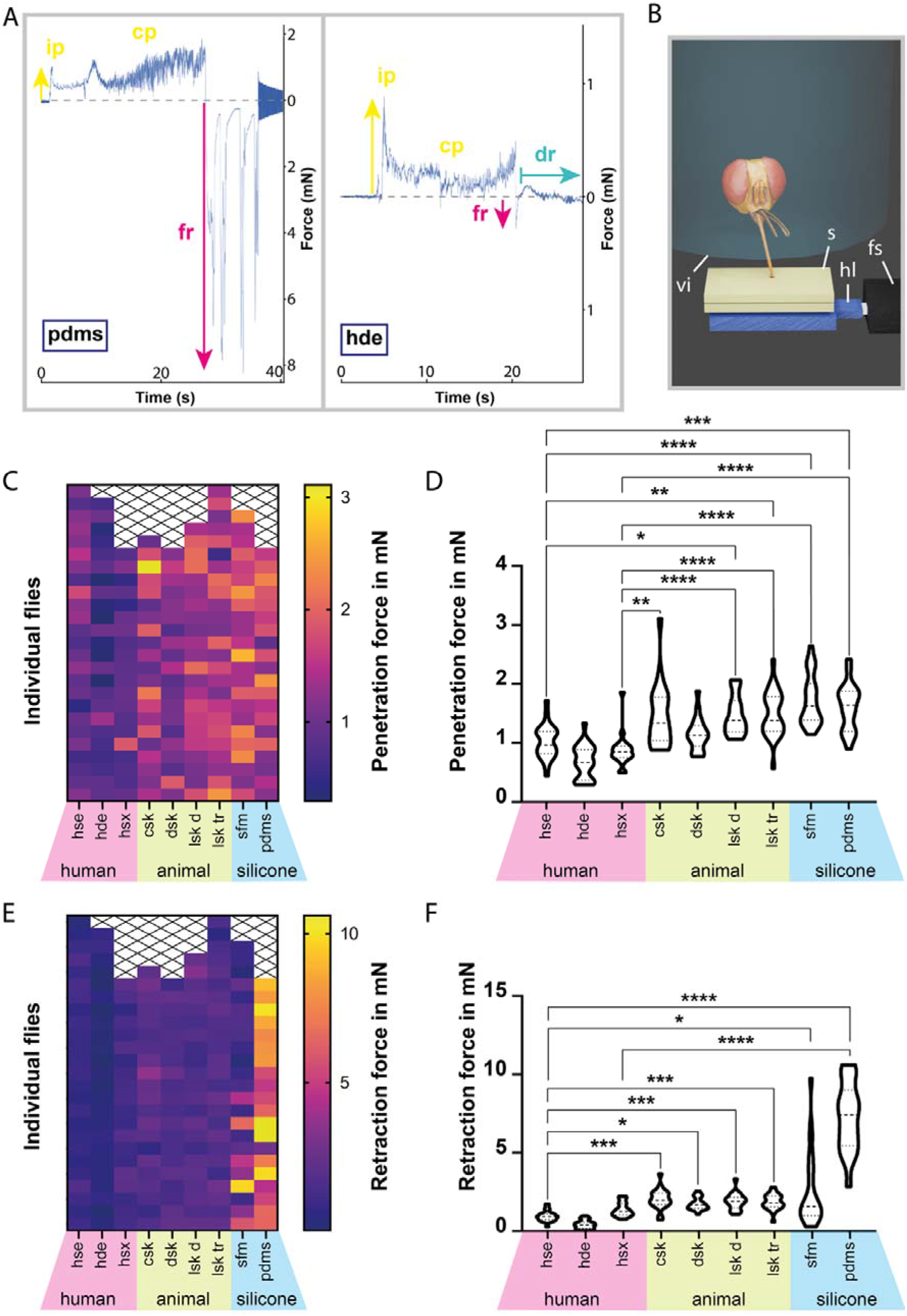
Force measurements during proboscis-substrate interaction. **A:** Example force plots of tsetse fly probing into PDMS and a hde, showing initial penetration, continued probing, and forced retraction. During probing on the hde, a drinking phase is visible. **B:** Visualization of the force measurement setup. A starved tsetse fly was held in a testing vial above a sample, which was mounted on a 3D-printed platform connected to a force sensor. The sample was heated to simulate host body temperature, triggering probing behaviour. **C, E:** Heatmaps of the penetration (C) and retraction (E) forces measured during fly probing. Each square represents a single probing event, grouped by substrate on the x-axis. **D, F:** Violin plots of the penetration (D) and retraction (F) forces measured during fly probing. The plots display full measurement ranges, with dashed lines marking medians and dotted lines marking quartiles. Statistical significance was determined using the Kruskal-Wallis test; *, p<0.05; **, p<0.01; ***, p<0.001; ****, p<0.0001. All values for penetration and retraction forces are provided in Supplementary material 2. **Abr**.: cp, continued probing; csk, cow skin; dr, drinking; dsk, deer skin; fr, forced retraction; fs, force sensor; hde, human dermal equivalent; hl, holder; hse, human skin equivalent; hsx, human skin explant; ip, initial probing; lsk d, lizard skin (dorsal); lsk tr, lizard skin (tailroot); s, sample; sfm, silicone feeding mat; sl, silicon layer; ts, tsetse fly; vi, testing vial.

While retraction generally marks the end of feeding, retractive force spikes also occurred during probing, when the proboscis was rhythmically pushed and retracted within the substrate (Figure 8A). Median retraction forces were highest for PDMS with 7.42 mN, and lowest for human skin samples, ranging from 0.38 mN to1.26 mN (Figure 8E, F). Animal skins exhibited median retraction forces of 1.97 mN for cow, 1.66 mN for deer, and 1.80 - 1.88 mN for lizard skins (Figure 8E, F). Statistical analysis confirmed that, overall, retraction forces significantly exceeded penetration forces (Figure S4A, B), even when silicone-based samples were excluded (Figure S4C, D).

### Analysis of tsetse fly blood meal parameters

In addition to force measurements, our experimental setup enabled real-time quantification of blood uptake parameters (Figure S5, Supplementary material 3), which were measured during probing on dermal equivalents. A characteristic drop in force signalled the onset of drinking (Figure 8A), reflecting the minimal proboscis movements that occur during this phase (Video 4). To calculate the ingested volume, we first determined the sample’s initial and final weights measured by the force sensor, yielding the total weight loss (Figure S5A). Evaporation was modelled as a linear function over the measurement period, and the evaporated weight was subtracted from the total weight loss to obtain the net drinking rate. Across 24 male flies, the average liquid uptake rate was 0.9 mg/s (Figure S5A), corresponding to a mean total intake of 26.2 mg per feeding event (Figure S5B). On average, each blood meal lasted 36.5 s (Figure S5C).

## Discussion

Currently, our understanding of the physical forces and biomechanical events involved in the majority of vector-host interactions remains limited, particularly regarding their role in host selection and feeding performance.

Attachment of the tsetse fly to the host surface is the first step in vector-host interaction. Hence, we measured the attachment of the tsetse fly to various surfaces (Figure 2, Video 1). Overall, attachment was strongest on glass or smooth epoxy resin, likely because smooth surfaces allow optimal engagement of the acanthae tips (Figure 2D).

However, when considering structured surfaces only, friction forces generally increased with roughness from 0.3 to 12 µm (Figure 2B, C, E, F). Attachment on the 9 µm surface appeared to be weaker than on 3 or 12 µm, possibly due to acantha spacing leading to partial contact loss at specific roughness’s (Figure 1C-H, Figure S1B, Figure 2D) (Kovalev et al., 2018).

In 2021, Büscher and colleagues reviewed attachment safety factors of 21 insect species, which ranged from 17 to 1130 across various experimental set ups, with a maximum of 146 for measurements on smooth surfaces. With average safety factors of ∼ 54 - 55 on glass (Figure 2E, F, H), the tsetse fly falls near the middle of this range. Its attachment performance is comparable to that of the Colorado potato beetle (*Leptinotarsa decemlineata*) with a safety factor around 60, as reported by Voigt and colleagues (2008). By contrast, non-blood-feeding hoverflies of the Syrphidae family show lower attachment performance, with safety factors of ∼ 25 - 30 on smooth polyvinyl chloride surfaces (Gorb et al., 2001).

At the high end of the spectrum, *Crataerina pallida* (Hippoboscidae) attaches to smooth surfaces with a safety factor of ∼ 90 (Petersen et al., 2018). These louse flies, close relatives of *Glossina* spp., are highly specialized (Hayer et al., 2022; Petersen et al., 2018) obligatory ectoparasites of mammals and birds. They are known to transmit trypanosomes in birds (Santolíková et al., 2022) and spend a significant portion of their lives on their respective hosts (Hayer et al., 2022; Petersen et al., 2007; Yatsuk et al., 2023). In contrast, tsetse flies do not face the same selective pressures for permanent host contact, and their attachment system thus represents a functional compromise: adequate for brief attachment during feeding or rest yet adapted for frequent flight. Therefore, with respect to attachment, tsetse flies function as generalists without specialized adaptations.

Once host contact is established, the tsetse fly can begin feeding using its multi-part proboscis. Like the flexible fascicle of mosquitoes (Martin-Martin et al., 2023), the proboscis of the tsetse fly exhibits the ability to reach in various directions beneath the skin (Reuter et al., 2023). This flexibility is crucial for expanding its probing area. In addition, the dorsoventral variation in stiffness at the base of the labellum (Figure 6) might be involved in guiding probing movements, analogous to the mechanism observed in the wasp ovipositor (Cerkvenik et al., 2019).

Furthermore, hollow segments within the labrum are thought to modulate haemolymph pressure, thereby dynamically regulating proboscis rigidity and influencing labellum movements, particularly eversion (Popham & Abdillahi, 1975).

The remarkably smooth wound margins produced by the proboscis in host tissue (Figure 7D-G) can be attributed to the cooperative action of its different tooth-like structures. Particularly the rasping teeth (Figure 5A, E, F, G, J, Figure S3B, C, and Video 3) function as fine abrasives that facilitate the breakdown of elastic tissue compartments. Furthermore, the secretion of saliva may act as lubricant, reducing friction not only between the mouthparts (Lehane, 2005), but also between the proboscis as a whole and host tissue. Collectively, these features decrease resistance, allowing the proboscis to move smoothly through the tissue.

Based on its structure and the orientation of the teeth (Figure 5), the fully opened labellum not only allows for blood uptake but might also serve as an anchor to secure the proboscis in place throughout the blood meal (Figure 7A-C). This mechanism may well enable the fly to resist host defences, such as the shake-off reflex.

When probing on lizard skin, we found that the fly avoids random penetration, instead probing specifically between large scales where the surface is composed of presumably softer regions (Figure 7M, N). This targeted probing likely reflects an adaptive strategy to overcome the physical barrier of reptilian skin and may similarly apply to other hosts with varying skin types.

To further investigate the interplay between the fly’s anatomy and the host’s skin properties, we measured the probing forces exerted on synthetic and natural substrates (Figure 8B). When interpreting absolute force magnitudes, it is important to bear in mind that our samples do not fully recapitulate physiological conditions. Skin explants and skin equivalents may behave differently to skin under active perfusion and native tissue tension, as may our fixed and frozen animal skin samples. Nevertheless, comparative force measurements revealed consistent biomechanical signatures across substrates, suggesting that the observed force patterns reflect fundamental aspects of the feeding mechanism that are likely relevant *in vivo*. The force patterns provided precise characterization of the distinct bloodmeal stages and revealed that throughout probing, penetration forces alternated with even stronger retraction forces (Figure 8). These intermittent pullbacks during probing, combined with the outward-facing teeth of the everted labellum, may cause primary tissue disruption. This could serve to enlarge the biting channel and improve access to deeper or more resilient blood vessels, thereby creating a larger blood pool for feeding.

Although visual observation of a tsetse bite gives the impression of extremely high forces (see Supplemental Video 1 in Reuter et al., 2023), our measurements revealed values in the low mN range. On human full-thickness skin equivalents, the tsetse fly exerted median bite forces of 0.97 mN (Figure 8C-F, Supplementary material 2). Nevertheless, this is ∼ 54 times greater than the average mosquito bite force on human skin with ∼ 18 µN (Kong & Wu, 2009). The mosquito’s penetration mechanism involves high-frequency fascicle vibration to reduce tissue resistance and minimize penetration forces (Kong & Wu, 2009). Oscillatory movements were also observed during tsetse probing (Video 4). Notably, these oscillations appeared predominantly during active blood uptake rather than during the initial penetration phase, suggesting that they are associated with ingestion rather than insertion. Whether they facilitate blood flow, prevent occlusion of the feeding canal, or simply reflect pump activity remains unknown. In general, the mosquitos feeding apparatus compares well to a highly flexible syringe designed for precise and delicate capillary feeding (Ramasubramanian et al., 2008). In contrast, the tsetse fly, as a blood pool feeder, resembles a flying saw, with a proboscis that is not only more forceful but also deliberately destructive.

Beyond recording probing forces, our setup also enabled quantification of blood uptake parameters (Figure S5). The bloodmeal volume of *G. morsitans* averaged 26.2 mg per feeding event, comparable to values reported for *G. austeni* in a previous study with an average intake of 26 mg (Margalit et al., 1972). In contrast, the mean feeding duration of *G. morsitans* was 36.5 s, markedly shorter than the 49 – 204 s reported by Margalit and colleagues for *G. austeni*. These differences in feeding time may reflect species-specific feeding behaviours but could also arise from methodological differences between experimental setups.

It is also important to note that our experiments were conducted using uninfected flies. Previous studies have shown that trypanosome infection can alter feeding behaviour, leading to increased probing activity and prolonged feeding times (Jenni et al., 1980; Van den Abbeele et al., 2010). These effects have primarily been attributed to infection-induced changes in saliva composition and the resulting interactions with host blood (Van den Abbeele et al., 2010). However, a different study found no significant effects of infection with either salivary gland-resident *T. brucei* or with proboscis-colonizing species such as *T. congolense* and *T. vivax* on Glossina feeding behaviour (Moloo, 1983). Furthermore, although infection-associated transcriptional changes in the salivary glands and proboscis have been reported (Awuoche et al., 2017; Matetovici et al., 2016), there is currently no direct evidence that trypanosome infection alters the mechanical properties or function of the mouthparts themselves.

Overall, our observations of trypanosomes within the fly’s hypopharynx, labial gutter, and host tissue are consistent with the established model of salivary-gland-derived parasite release during probing and feeding. Their presence beyond the immediate feeding canal is consistent with rapid local dispersal following inoculation, as described previously (Reuter et al., 2023). This process may be facilitated by the extensive tissue disruption caused by the tsetse mouthparts, although this hypothesis will require direct experimental testing.

Ultimately, the objective of this study was to investigate how tsetse flies can feed on a seemingly random selection of animals with highly diverse skin structures. In our detailed anatomical studies and force measurements, we surprisingly have not found a major single trait that explains the fly’s feeding versatility.

Instead, our results indicate that this capability emerges from the combined effect of multiple, more subtle traits. In particular, the proboscis generates broadly similar penetration forces across a wide range of skin types, suggesting a generalised mechanical mechanism rather than host-specific optimisation. The intricate architecture of the labellum and the strong retractile forces during probing likely contribute to efficient penetration and blood pool formation across heterogeneous substrates. Behaviourally, tsetse flies further increase feeding success by flexibly targeting mechanically favourable sites, such as the softer interscale regions on lizard skin, rather than relying on specialised morphological adaptations.

This composite strategy likely reflects evolutionary fine-tuning that enables the remarkably broad host range of tsetse flies. Such feeding versatility provides an ecological framework that has undoubtedly contributed to the extraordinary evolutionary success of African trypanosomes, which parasitise virtually all warm-blooded sub-Saharan vertebrate species.

## Methods

### Fly keeping

Tsetse flies were kept in Roubaud cages at 27 °C with 70 % humidity. They were fed three times per week with 37 °C defibrinated sheep blood provided on a metal tray through a silicone feeding mat (Feeding membrane, Slovak University of Technology, Bratislava, Slovakia).

### Fly infection

For tsetse fly infections, the pleomorphic *Trypanosoma brucei brucei* strain EATRO 1125, serodeme Antat 1.1 was used (Le Ray et al., 1977). Culture of slender bloodstream forms, stumpy differentiation, and infection of flies with stumpy forms were performed as described in Reuter et al., 2023.

### Fly dissection

Flies were sedated with chloroform in a glass vial, and wings and legs were removed. Each fly was transferred into 50 μl PBS on a glass slide, where the region of interest was dissected using fine forceps under a stereomicroscope (Wild Heerbrugg, Heerbrugg, Switzerland).

### Confocal laser scanning microscopy (CLSM)

The material distribution of the cuticle was analysed using a confocal laser scanning microscopy (CLSM)-based method which was established by Michels and Gorb (Michels & Gorb, 2012) and validated previously (Peisker et al., 2013).

Fly samples preserved in 70 % ethanol were soaked in distilled water, transferred to glycerol (≥ 99.5 %, free of water, Carl Roth GmbH and Co. KG, Karlsruhe, Germany), and mounted in glycerol on glass slides. To prevent sample damage, adhesive reinforcement rings were placed between slide and coverslip (Michels & Büntzow, 2010).

Imaging was performed using a Zeiss LSM 700 confocal laser scanning microscope (CLSM) (Carl Zeiss Microscopy GmbH, Jena, Germany), equipped with 405 nm, 488 nm, 555 nm, and 639 nm stable solid-state lasers. Autofluorescence signals of the insect cuticle were detected using a bandpass emission filter (420–480 nm) and long-pass emission filters transmitting wavelengths ≥ 490 nm, ≥ 560 nm or ≥ 640 nm, respectively (Büsse and Gorb, 2018). Imaging was performed with 10 × (Zeiss EC Plan-Neofluar, NA = 0.45) and 20 × (Zeiss Plan-Apochromat, NA = 0.8) air immersion lenses. Autofluorescence micrographs were pseudo-coloured (each at 50 % saturation) based on the method by Michels and Gorb (2012) and combined using maximum intensity projection in ZEN software (Zeiss Efficient Navigation; Carl Zeiss MicroImaging GmbH). The resulting colour gradient indicates cuticle composition: blue = resilin-rich (soft), green = moderately sclerotized, red = highly sclerotized (rigid).

### Conventional sample processing for scanning electron microscopy (SEM)

Different fixation and dehydration approaches were applied. To preserve external morphology, fly parts were fixed in 70 % ethanol for 30 min at room temperature (RT) and immediately dehydrated through an ascending ethanol series (80 %, 90 %, 96 %, and 2 x 100 %), followed by 3 x 20 min washes in 100 % acetone. For the proboscis fixed during probing on the feeding mat, flies were frozen mid-penetration by immersing the sample in 100 % ethanol precooled to −80 °C (Peinert et al., 2016). Samples were stored at −80 °C in ethanol for two weeks, followed by 3 x 20 min ethanol exchanges. Ethanol was replaced with 100 % acetone, which was changed 2 x 20 min.

For regular SEM imaging, samples were fixed in either Karnovsky fixative (2 % PFA, 2.5 % glutaraldehyde in 0.1 M cacodylate buffer, pH 7.4) (Karnovsky, 1964) or 6.25 % glutaraldehyde in 0.075 M Sörensen buffer (pH 7.4) overnight at 4 °C. After removing the fixative, samples were washed 5 x 3 min in cold buffer (0.05 M cacodylate or 0,075 M Sörensen), incubated in 2 % tannic acid and 4.2 % sucrose in 0.05 M cacodylate buffer (pH 7.4) for 1 h at 4 °C, washed 3 x 5 min with 4 °C cold ddH_2_O, and dehydrated in ascending cold acetone (30 %, 50 %, 70 %, 80 %, 90 %, 96 %, and 2 x 100 %, each for 30 min), followed by 3 x 30 min incubation in 100 % acetone.

Critical point drying was performed using a Bal Tec 030 critical point dryer (Bal-tec™, Los Angeles, USA) and samples were placed in a sputter coater (Bal-tec SCD 050) for 150 s with 10 nm gold/palladium.

### Resin-embedded sample processing for SEM

The cow skin sample with inserted proboscis was fixed with 2.5 % glutaraldehyde in 0.05 M cacodylate buffer, pH 7.2, overnight at 4 °C. The sample was washed 5 x for 3 min with precooled 0.05 M cacodylate buffer and postfixed in OsO_4_ in 0.05 M cacodylate buffer, pH 7.2, for at least 90 min at 4 °C. OsO_4_ was removed by 5 x 3 min washes with cold ddH□O. Afterwards, the sample was stained with 0,5 % uranyl acetate overnight at 4 °C. Followed by 3 x 5 min washes with cold ddH□O. Samples were dehydrated using ascending concentrations of cold acetone (30 %, 50 %, 70 %, 80 %, 90 %, 96 %, and 2 x 100 %, each for 30 min), followed by final dehydration with propylene oxide for 5 x 3 min and transfer to a 1 : 1 mixture of epoxy resin (Epon) and propylene oxide for several hours at RT. Samples were immersed in 3 x 100 % Epon, 2 x 2 h, followed by a final incubation until the samples sunk to the bottom of the silicon form. Samples were placed in a preheated oven and polymerised at 60 °C for 48 h and the Epon block was trimmed using a razor blade. Sections of 500 nm were cut with a diamond knife (Jumbo diamond, DiATOME, Nidau, Switzerland) and transferred to polylysine slides. The sections were contrasted with 2 % uranyl acetate in H□O for 10 min and washed 3 x with ddH□O. H_2_O was pre-boiled to reduce CO_2_ and cooled to RT, preventing precipitation during subsequent staining (Reynolds, 1963). Sections were incubated with 50 % Reynolds lead citrate prepared in pre-boiled ddH□O for 5 min, followed by two washes and a final rinse with pre-boiled ddH□O. Sections were dried with compressed air, and excess glass around the sections was removed. The remaining glass with sections was mounted onto electron microscopy stubs using conductive tape, and conductive silver was applied to the edges and blank areas to reduce charging. Samples were then sputter-coated with a 5 nm layer of carbon (CCU-010, Safematic GmbH, Switzerland).

### SEM imaging

SEM imaging was performed using a JEOL JSM-7500F microscope (Jeol, Tokyo, Japan). Complete fly parts or larger skin samples were imaged using a lower secondary electron detector (LEI), while serial sections were imaged with a low-angle backscatter electron (LABE) detector, each operated at 5 kV.

### Micro-computed tomography (µCT)

The tsetse head was fixed in 6.25 % glutaraldehyde in 0.05 M cacodylate buffer (pH 7.2) overnight at 4° C, then washed 3 x 5□min in ddH□O. Samples underwent dehydration in ascending ethanol (50 %, 70 %, 80 %, 90 %, and 96 % for 20 min each, and 2 x 30 min at 100 %) and were transferred to 100 % acetone for 3 x 15 min each. To ensure maximum dehydration, the acetone was refreshed once more prior to critical point drying using a Bal Tec 030 critical point dryer (Bal-tec™, Los Angeles, USA). µCT data was acquired using a SkyScan 1172 (Bruker micro-CT, Kontich, Belgium) tabletop µCT.

### Focused Ion Beam Scanning Electron Microscopy (FIB SEM)

The proboscis of an infected tsetse fly was high-pressure frozen using an EM HPM100 machine (Leica, Wetzlar Germany) and automatic freeze substitution was performed as described by Stigloher et al. (2011). In short, the frozen sample was transferred from liquid nitrogen into an EM AFS2 freeze substitution system (Leica Microsystems, Germany) and incubated with 0.1 % tannic acid and 0.5 % glutaraldehyde in anhydrous acetone at -90 °C for 96 h. The samples were washed with anhydrous acetone for 4 x 1 h at -90 °C, then contrasted with 2 % OsO_4_ in anhydrous acetone at -90 °C for 28 h. Temperature was raised to −20□ °C over 14□ h, held for 16□ h, then increased to 4□ °C over 4 □h. Samples were washed 4 × 30□ min with acetone at 4□ °C and the temperature was raised to 20 °C within 1 h. The samples were transferred to a 1 : 1 mix of acetone and Durcupan™ ACM (Sigma-Aldrich, Merck Group, St. Louis, USA) and stored for 5 h at RT to facilitate infiltration (Schieber et al., 2017). The old resin was replaced sequentially with 90 % durcupan in acetone overnight at 4 °C, followed by 3 x 2 h 100 % durcupan. Residual resin was removed by gently moving the sample over Aclar film with a wooden toothpick. The portion of Aclar with the sample was cut out and cured at 60 °C for 48 h, then stored in a dark, dry place (Schieber et al., 2017).

The sample was mounted onto SEM stubs and sputter coated with 10 nm Tungsten to dissipate charges (ACE 600, Leica, Wetzlar, Germany). In the Crossbeam 540 (Zeiss, Oberkochen, Germany), an 800 nm platinum layer was deposited on the sample surface, and a cross-section was opened with a 15 nA ion beam and polished with 7 nA. Data were acquired using Atlas 5 (Zeiss, Oberkochen, Germany) at 1.5 kV (1200 pA) with the EsB detector (450 V) and FIB at 3 nA in continuous mode, producing a voxel size of 15 x 15 x 50 nm. Post-processing in Fiji (Schindelin et al., 2012) included SIFT alignment, cropping, inversion, Gaussian blur (2/1), local contrast enhancement (CLAHE: 127, 256, 2) and 2 x binning in x/y.

### Image processing

3D reconstruction and visualization of the volume and isosurface renderings of the µCT and FIB-SEM data was done with Amira 6.4 (Thermo Fisher Scientific, Massachusetts, USA). Amira surface files (.iv) of the tsetse head model were converted to object files (.obj) with Transform2 64 bit (Heiko Stark, Jena, Germany). For refinement and rendering, 3D models were processed with Autodesk Maya 2023.2 (Autodesk, San Francisco, USA) and Blender 3.5.0 (Stichting Blender Foundation, Amsterdam, Netherlands).

Image plates, illustrations and measurements were created using Adobe photoshop 2020 (Adobe Inc., San José, USA), Adobe illustrator 2020 (Adobe Inc., San José, USA) and Fiji (ImageJ 1.53t, National Institutes of Health, USA).

### Macro photography and videography

Macroscopic images and force measurements were recorded by a Canon EOS 80D DSLR camera (Canon, Tokyo, Japan) equipped with a Canon EF 100 mm f2.8L Macro IS USM lens (Canon, Tokyo, Japan). Focus stacking of image series was performed using Helicon Focus (Helicon Soft Ltd, Kharkiv, Ukraine) and Zerene stacker version 1.04 (Zerene Systems LLC, Richland, USA). Video 1 was taken with a Samsung Galaxy S6 (Samsung, Seoul, South Korea) and Video 4 with the Canon EOS 80D DSLR camera equipped with a Canon EFS 18-135 mm objective (Canon, Tokyo, Japan).

### Friction force measurement

Friction forces were measured following the method of Gorb et al. (2001). The setup consisted of a motor-driven rotating drum fitted with discs of varying materials and surface roughness. A fiber-optic sensor and laser beam detected the moment the fly lost contact with the disc. Simultaneously, the setup, controlled via computer and software (Throw the fly 1.0, TETRA GmbH, Ilmenau, Germany), provided precise measurements of multiple parameters at the moment of contact loss, including motor speed (revs min^-1^), sensor displacement from the rotor center d (cm), duration of rotation Δt (s), and time between sensor signals Δt1 (s). Prior to the experiment, all flies were starved and weighed, and their wings were removed. Paired groups (n=10) of male and female flies were used. Each fly was individually placed on the midsection of the disc, after which the position was verified.

Friction forces were quantified by assuming detachment occurred when the centrifugal force exceeded the maximum static friction. Following Gorb et al. (Gorb et al., 2001), the friction force is calculated as: F_friction_ = m × r × ω²; where m = mass of the fly (kg), r = radial distance from the centre of rotation (m), ω = angular velocity (rad/s). The angular velocity is derived from the rotation frequency f using: ω = 2 × π × f. For example, a fly with a mass of 2 mg positioned 3 cm from the centre and exposed to a rotation frequency of 10 Hz has an angular velocity of: ω = 2 × π × 10 = 62.83 rad/s This results in F_friction_ = 0.000002 × 0.03 × (62.83)² ≈ 237 µN.

The Friedman test was used to assess differences in friction forces and safety factors between surfaces, separately for male and female flies. To compare overall attachment capabilities (forces and safety factors) between male and female flies across all surfaces, the Mann–Whitney test was applied.

### Proboscis force measurement

The measurement setup was custom designed. A 3D-printed sample holder (Prusa Research s.r.o., Prague, Czech Republic) was attached to a FORT10g Force transducer (World Precision Instruments, Inc., Sarasota, USA) connected to a BIOPAC system (BIOPAC Systems, Inc., USA), allowing detection of force signals and their conversion into electrical resistance. The samples were resized to fit the sample holder using razor blades and scalpels. Apart from the silicone samples, all tissue samples were placed in either PBS (animal samples) or defined media (human skin explants and skin equivalents).

An individual fly was placed in a 15 ml tube modified with a net at one opening and a pushing device at the other. The fly at the net opening was oriented approximately 1 mm above the sample to allow for proboscis-sample-interactions while preventing interference from the rest of the tsetse body. To stimulate probing, flies were starved for at least 24 h prior to the experiment and a heat source was placed close to the sample holder to imitate the body temperature of a vertebrate host. The temperature was monitored using an infrared camera (IC 080 V; Trotec, Heinsberg, Germany) and adjusted to avoid overheating and sample drying. The measurements were recorded and visualized with the software AcqKnowledge (BIOPAC Systems, Inc., USA). For each substrate, 20-25 flies were measured.

To compare proboscis penetration or retraction forces between different substrates, the Kruska-Wallis test was used. To generally compare penetration with retraction forces across all substrates or specifically across all animal samples, the One-sample Wilcoxon test was used.

### Statistical analysis

Statistical analyses were performed with Microsoft Excel 2021 (Microsoft, Washington, USA) and Prism 9.0 (GraphPad Software, Boston, USA).

### Origin of probing substrates for microscopy and proboscis force measurements

NativeSkin® skin explants (Ref. NSA08; Lot 20211025.02) from the abdominal region of a 36-year-old female with fair, normal skin (Fitzpatrick type 2) were obtained from Genoskin (Toulouse, France). Samples were embedded in a gel substrate to maintain hydration and viability during shipping and used for SEM imaging of proboscis penetration. Human skin explants were collected from surplus surgical tissue and stored in PBS. Excess fat was removed before use in proboscis force measurements.

Human dermal and full-thickness skin equivalents were prepared and cultured according to Reuter et al., 2023.

Deer skin (*Capreolus capreolus)* from a young animal was kindly provided by a hunter from Frankonia, southern Germany. Cow skin was provided by a farmer in southern Germany. Prior to experiments, the skin samples were depilated using disposable razors and cut into appropriately sized pieces with scalpels and surgical scissors. Samples were stored at −20 °C until use.

Skin from the middorsal region and tail root of the monitor lizard *Varanus niloticus* (Linaeus 1766; Inventory number: ZMFK 84206) was provided by Wolfgang Böhme and Thore Koppetsch (Museum Koenig in Bonn, Germany) and stored in 70 % Ethanol.

The feeding mat was obtained from the Slovak University of Technology, Bratislava, Slovakia.

For the polydimethylsiloxane (PDMS) substrate, PDMS (Sylgard 184) (Sigma-Aldrich, Merck Group, St. Louis, USA) was mixed in a 10:1 ratio with a curing agent, poured into a plastic dish and subsequently cured at RT for 24 h.

### Ethical clearance statement

Human skin explants originated from excess skin of a surgical procedure. Male donors aged 2 - 5 years contributed normal primary human epidermal keratinocytes (NHEK) and dermal fibroblast (NHDF) via preputial skin biopsies. Fully informed consent was obtained from the donor’s legal representatives. Approval of the local ethical board of the University of Würzburg (vote 182/10 and 280/18-SC) was given.

## Acknowledgements

We are grateful to Georg Krohne, who provided initial SEM micrographs of the tsetse head, proboscis and tarsal features as well as helpful advice to electron microscopy techniques. We thank Jaime Lieb for the helpful discussions and critical reading of the manuscript. We thank Alyssa Bergmann Borges for providing essential statistical advice. We also thank Wiebke Möbius (Max-Planck-Institute, Göttingen, Germany) for giving us the opportunity to use her laboratory and the FIB-SEM. We are grateful to Hans Pohl (University of Jena, Germany) for constructive comments regarding freeze fixation for the probing assay of the fly proboscis, and Sebastian Markert (Hochschule für Technik und Wirtschaft des Saarlandes, Saarbrücken, Germany) for advice regarding the handling of samples for minimal resin embedding. We would like to acknowledge the staff of the Microscopy Core Facility of the Biocentre and Beate Vogt for their help in preparing the samples for electron microscopy. We express our gratitude to Christa Albert for helping with the skin explants. We thank Thore Koppetsch (Zoologisches Forschungsmuseum Alexander Koenig, Bonn, Germany) for help with the delivery of the monitor lizard sample. We also want to thank Matthias Leippe (Kiel University, Germany) for providing infrastructure for skin model maintenance during the measuring experiments and Heidrun Ließegang and Christine Blurton (Kiel University, Germany) for the technical assistance in the cell culture. We thank Matthias Peindl for providing the deer skin and Bianca Link for the cow skin used in the probing force measurements. We thank Christian Reuter for his contribution to recording the tsetse fly feeding on human skin models. Finally, we acknowledge support by Mohsen Jafarpour (Kiel University, Germany) in 3d design and printing of experimental setup parts.

## Supplementary material

**Video 1: Experimental setup for friction force measurements.** Shown are the rotating drum with a mounted disc and a positioned insect, surrounded by a protective enclosure made of soft foam material. The disc accelerates until the fly loses contact. This is the time point when the when the centrifugal force exceeds the maximum static friction. The friction forces and safety factors were calculated according to the equations from Gorb et al., 2001.

**Video 2: µCT scan of a tsetse fly head.** The left panel shows a sequential display of the acquired 2D µCT projection slices, while the right panel presents a 3D reconstruction of the complete dataset. Imaging was performed on a SkyScan 1172 tabletop µCT system.

**Video 3: FIB-SEM of the tsetse labellum.** Sequential cross sections reveal internal ultrastructure progressing from near the tip of the labellum downward. Data were acquired on a Crossbeam 540 (Zeiss) with the EsB detector in continuous milling mode. Reconstructed images from this dataset are shown in Figure 5A and Supplementary Figure 3B and C.

**Video 4: A tsetse fly taking a blood meal from human skin equivalents placed on sheep blood.** The skin equivalents were stacked to provide sufficient biting depth for the fly. Vibratory movements of the fly’s maxillae can be observed during feeding. The video was recorded in collaboration with Christian Reuter.

**Supplementary material 1: Tarsal attachment performance of tsetse flies expressed in friction force and safety factor.** Flies were placed on a motor-driven rotating drum fitted with discs of varying materials and surface roughness. A fiber-optic sensor with a laser beam detected the moment each fly lost contact during acceleration. Measurements were obtained from 10 female and 10 male flies, tested on six different surfaces. The table includes raw measurements from each trial, calculated friction forces and safety factors, and mean values per fly (based on 3–14 replicates). Friction forces and safety factors were calculated as described by Gorb et al. (2001). The safety factor, defined as the total friction force divided by weight force (mg, where m is body mass and g is gravitational acceleration), is a dimensionless quantity that enables cross-individual comparison.

**Supplementary material 2: Forces during proboscis-substrate interaction.** Starved tsetse flies were individually positioned above samples mounted on a 3D-printed platform connected to a force sensor and probing behaviour was triggered. Penetration and retraction forces of the proboscis were recorded by the sensor during probing on human, animal, and silicone substrates. The table provides forces measured for each fly, with 20–25 flies tested per substrate, along with mean and median values per substrate.

**Supplementary material 3: Liquid uptake parameters during tsetse feeding.** Parameters were recorded from 24 male tsetse flies feeding on dermal equivalents during proboscis force measurements. The table lists individual values for sample weight loss, evaporation-corrected drinking rate, blood meal weight, and feeding duration, along with mean and median values. Sample weight loss was determined by weight measurements from the force sensor; evaporative loss was estimated and subtracted to calculate the actual drinking rate.

---

**Figure Supplement 1:**
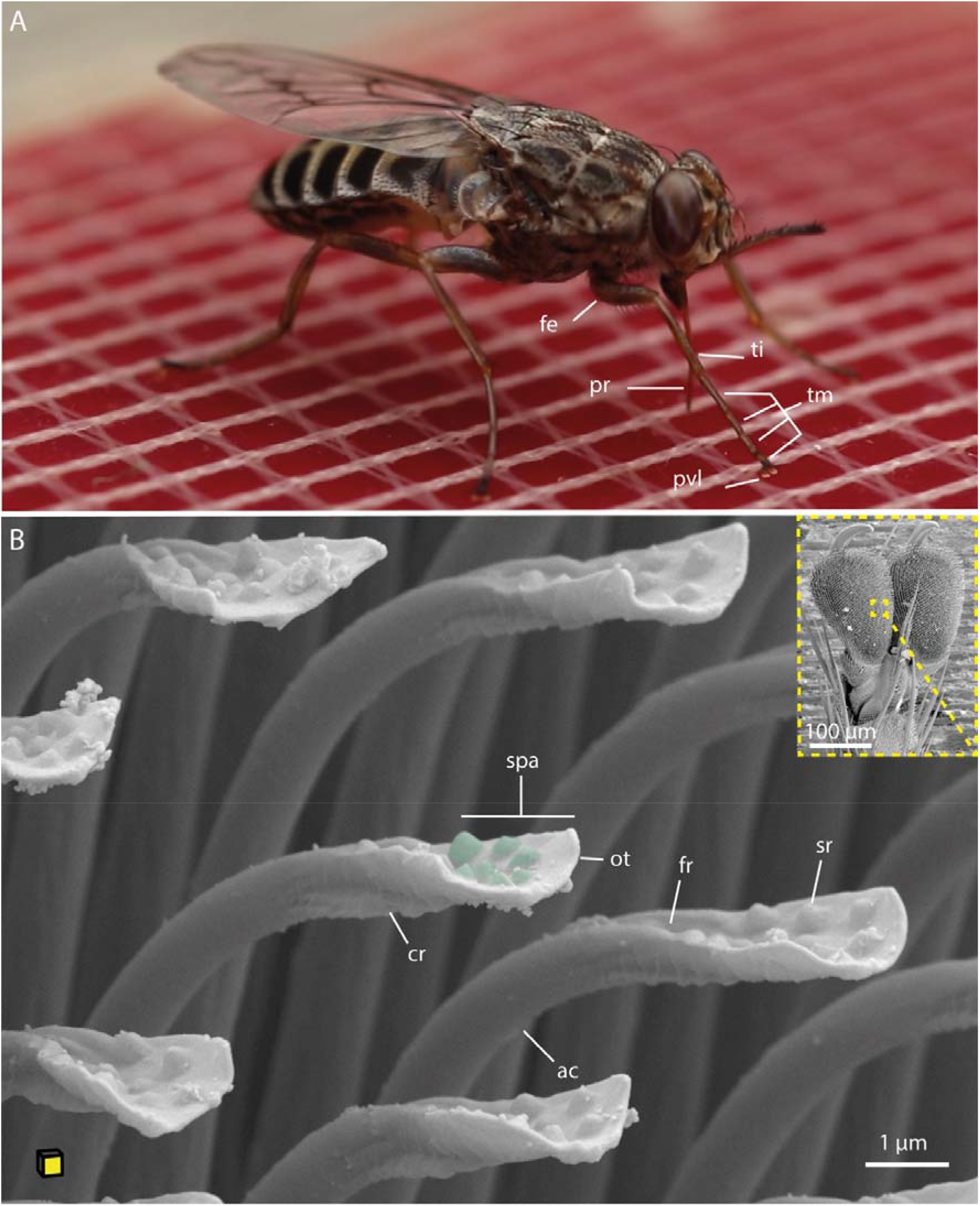
Blood-feeding tsetse fly and SEM images of tsetse tarsal adhesive structures. **A:** A tsetse fly taking a blood meal on a silicone feeding mat. **B:** Ventro-lateral overview of the pulvillus with a magnified view of the acanthae, showing fine crests on the underside and knob-like surface modifications (green) on the upper side. The yellow plane of the cube indicates the orientation of the tarsus attachment surface. **Abr.:** ac, acanthae; cr, crest; fe, femur; fr, furrow; ot, oblate tip; pr, proboscis; pvl, pulvillus; spa, spatula; ti, tibia; tm, tarsomeres.

**Figure Supplement 2:**
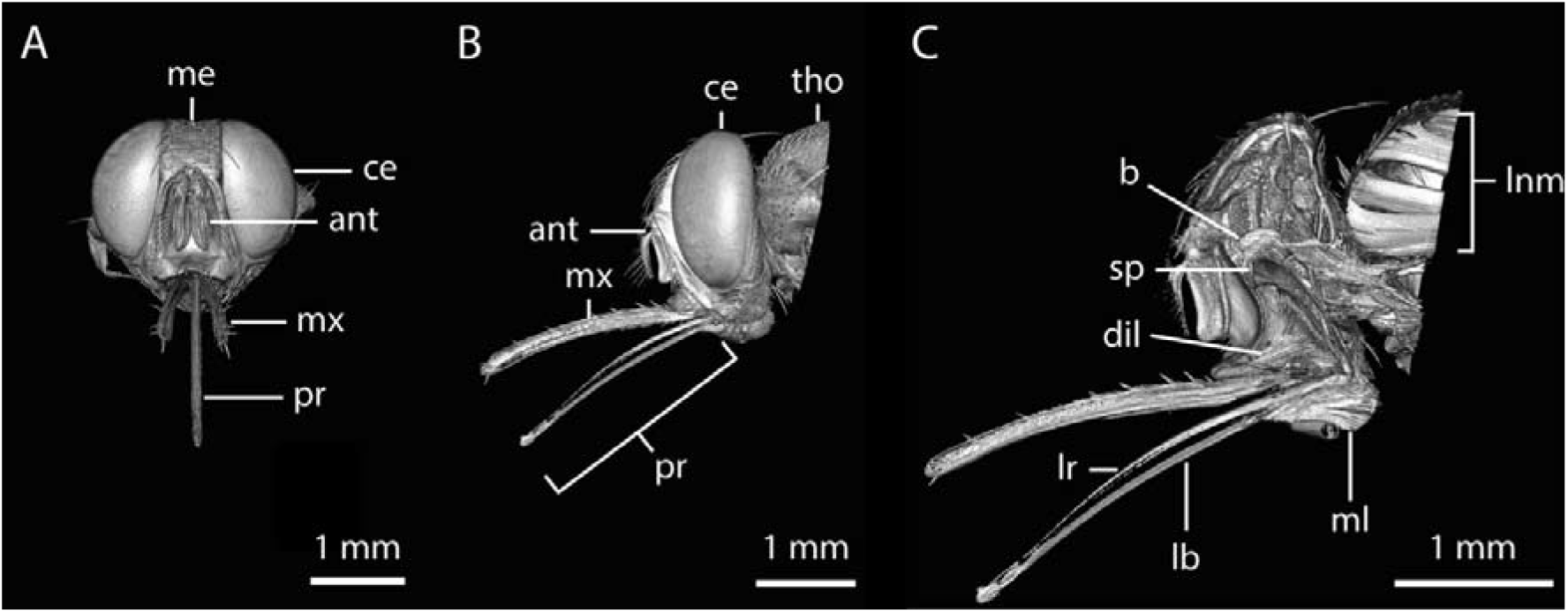
Rendering of a tsetse head derived from micro-computed tomography (µCT) data. Related to Figure 3. **A:** Frontal view. Video 2 provided with this manuscript shows a 360° rotation of this 3D rendering. **B:** Lateral view. **C:** Lateral view of the digitally generated cross section. **Abr.:** ant, antenna; b, brain; ce, compound eye; dil, pharyngeal dilator muscles; lb, labium; lr, labrum; lnm, longitudinal muscles; me, median eye; ml; labial muscles; mx, maxillary palp; pr, proboscis; sp, suction pump; tho, thorax.

**Figure Supplement 3:**
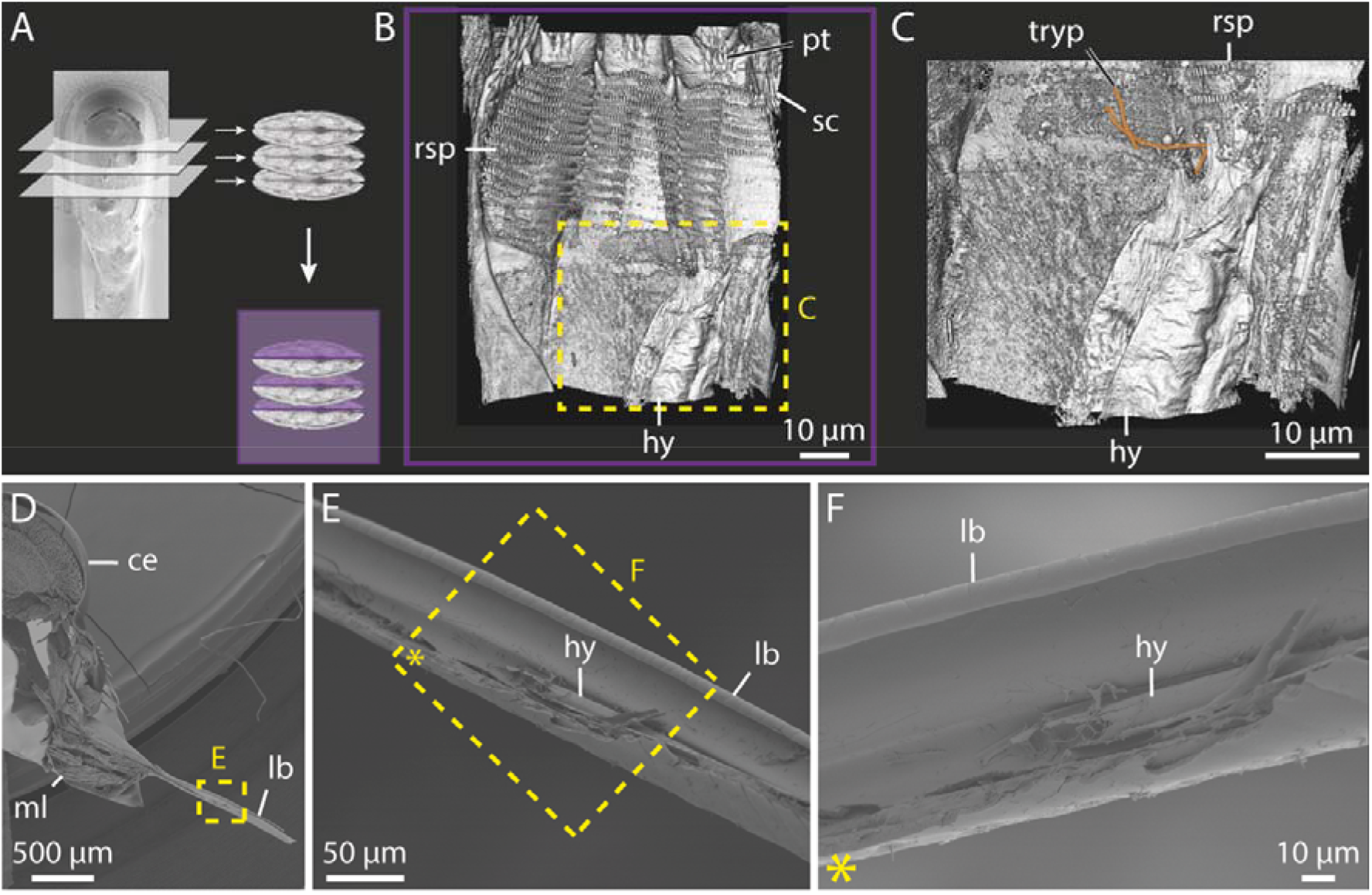
FIB-SEM imaging of trypanosome-infected and uninfected tsetse labella and SEM imaging of the dissected proboscis. Related to Figure 5. **A:** The labellum was imaged gradually layer by layer from the tip using FIB-SEM (Video 3), reconstructed in 3D, and a ventral view of the interior was generated, as indicated by the purple plane. **B:** Ventral overview of the inside of the digitally dissected closed labellum. **C:** Magnified view of B, trypanosomes are highlighted in orange. **D:** Lateral overview of a dissected head and proboscis, exposing labial muscles in the bulbus. **E:** Close-up of C, showing parts of the dissected labium. **F:** Magnified and rotated view of D, showing the labial gutter and part of the hypopharynx. **Abr**.: ce, compound eye; hy, hypopharynx; lb, labium; ml, labial muscles; pt, prestomal tooth; rsp, rasping teeth; sc, scales; tryp, trypanosomes.

**Figure Supplement 4:**
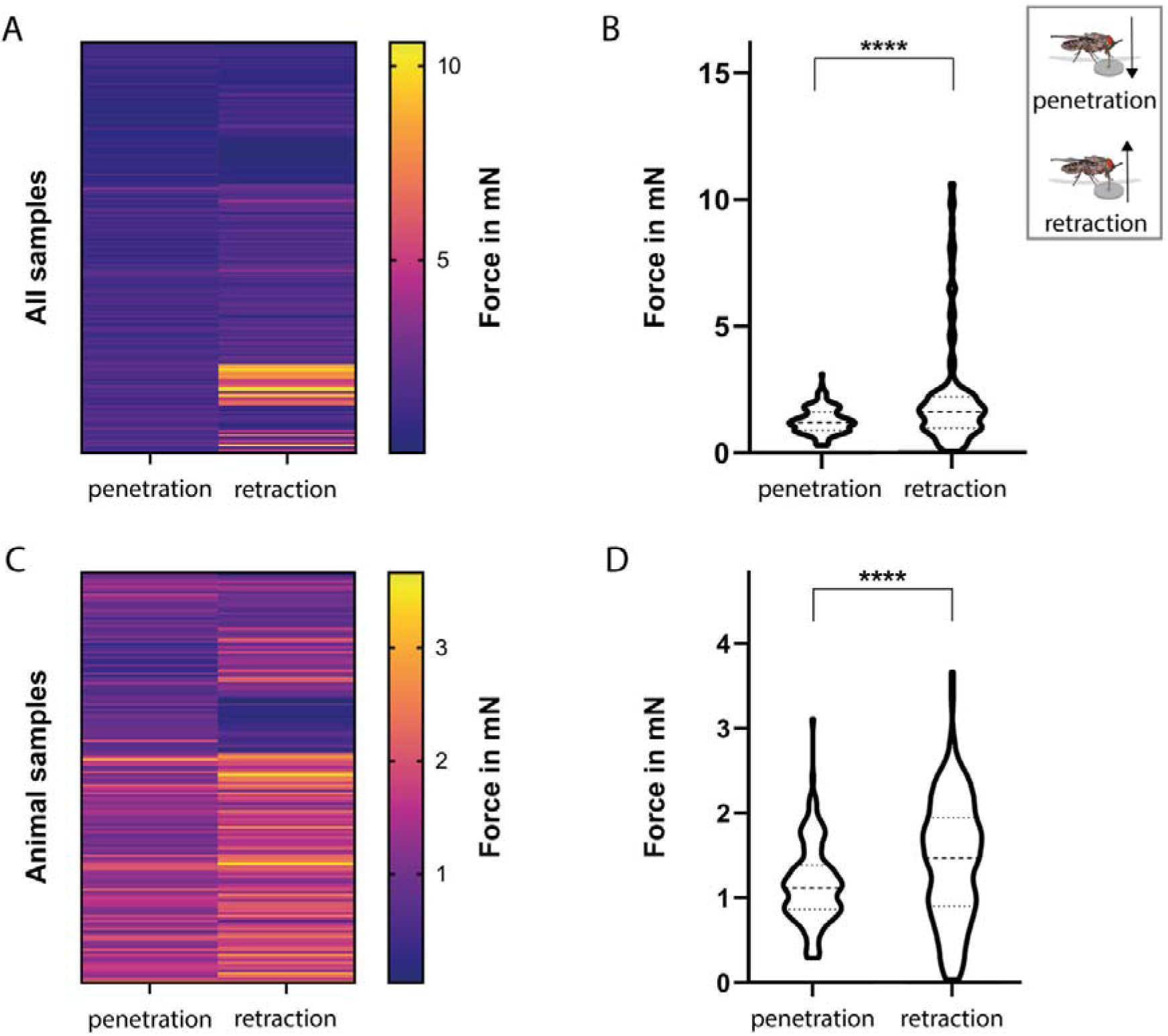
Comparison of penetration and retraction forces during proboscis-substrate interaction. Related to Figure 8. **A-B:** Heatmap and violin plot of penetration and retraction forces across all substrate types. **C-D:** Heatmap and violin plot of penetration and retraction forces across all animal samples (excluded are silicon and PDMS substrates). Statistical significance was determined using the One-sample Wilcoxon test; ****, p<0.0001.

**Figure Supplement 5:**
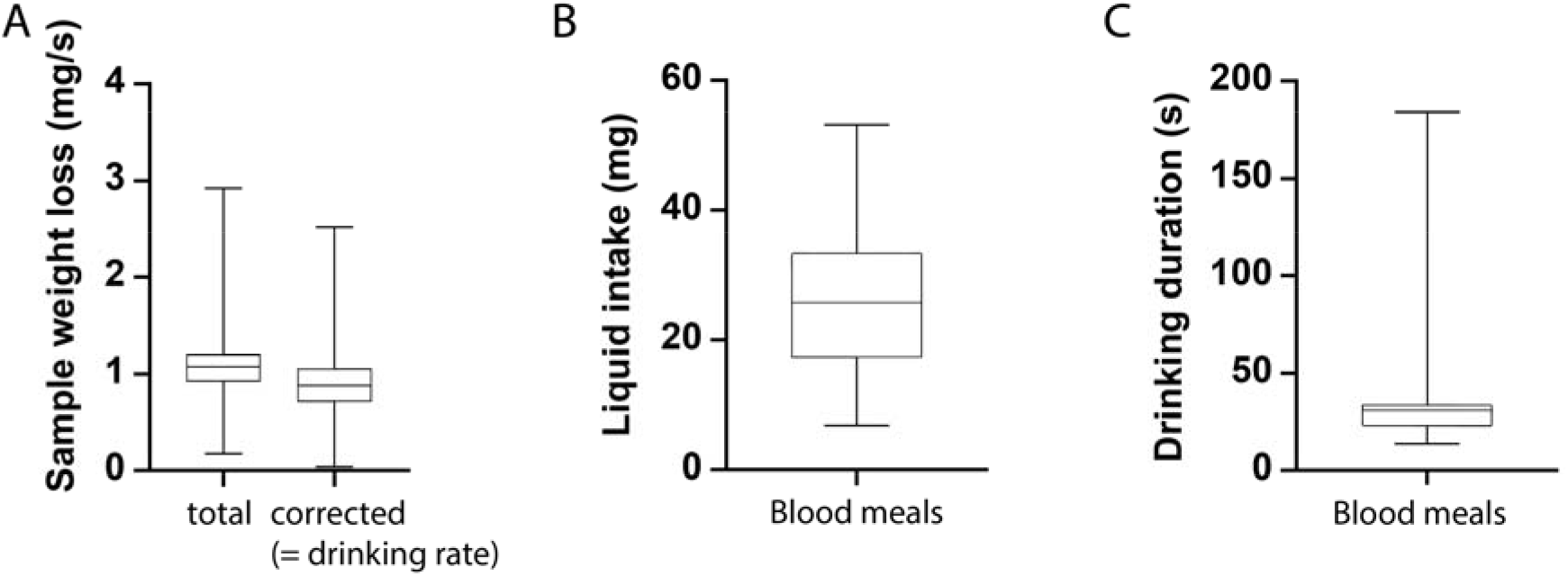
Quantification of liquid uptake dynamics during tsetse feeding. All parameters were determined from flies feeding on dermal equivalents during proboscis force measurements. Shown are boxplots of the sample weight loss over time (**A**), the blood meal weight (**B**), and the blood meal duration (**C**) for 24 tsetse flies. Boxes span the interquartile range, whiskers the full range, and a line marks the median. In A, “total” indicates the overall sample weight loss during the experiment, derived from weight measurements by the force sensor. Evaporative loss was estimated and subtracted, yielding the rate of liquid uptake (“drinking”) by the tsetse flies. All values are provided in Supplementary material 3.

